# Fruitful female fecundity after feeding *Gryllodes sigillatus* royal jelly

**DOI:** 10.1101/2022.05.17.492327

**Authors:** MJ Muzzatti, E McConnell, S Neave, HA MacMillan, SM Bertram

## Abstract

Dietary honey bee royal jelly increases insect growth rates and adult body size. Royal jelly could enhance mass insect production as a dietary supplement, but it is costly to produce. The mechanisms underlying the effects of royal jelly on growth remain unclear, and so it is valuable to understand the effects of royal jelly on a mass reared model species to try and augment body size in a more cost-effective manner. To determine the effect of royal jelly on a cricket species (*Gryllodes sigillatus*) farmed on mass for human consumption, we ran two experiments. In one experiment we tested the dose-dependent response of *Gryllodes sigillatus* to royal jelly using a range of diets across 0-30% w/w royal jelly. In another experiment we measured the individual-level life history responses of *Gryllodes sigillatus* to royal jelly over time by individually rearing freshly-hatched *Gryllodes sigillatus* on two separate diets: half were fed a commercially available cricket diet, while the other half were fed the same diet mixed with 15% w/w fresh royal jelly. Body size and mass measurements were recorded weekly for five weeks. We found the effects of royal jelly to be sex-dependent within crickets: females fed the royal jelly diet grew to be 30% heavier, and this effect was driven by significantly longer abdomens containing 67% more eggs compared to those fed the basal diet. There was a higher probability of crickets reaching adulthood after 35 days when fed royal jelly, and female mass was optimised at approximately 17% w/w royal jelly. Our results reveal that while a royal jelly dietary supplement can increase the yield of mass-reared insects, the life-history responses are species- and sex-specific.

## 1.0 Introduction

Honey bee royal jelly is a nutrient-rich substance produced and secreted by the hypopharyngeal and mandibular glands of nurse bees that has strong epigenetic influence on the polyphenism of caste development. Honey bee larvae fed royal jelly *ad libitum* will develop into queens, resulting in a caste with remarkable life history differences compared to other caste members including larger bodies, longer lifespan, and the ability to lay fertilized eggs (Haydak 1943; Ramanathan et al. 2018; Slater et al. 2020). Over the past ten years, the life history altering effects of royal jelly have been discovered to be interspecific, eliciting changes in body size, feeding, longevity, development time, and fertility in the fruit fly *Drosophila melanogaster* (Arruda et al. 2017; Gardner 1948; Kamakura 2011; Kayashima et al. 2012; Shorter et al. 2015), the nematode *Caenorhabditis elegans* (Detienne et al. 2014; Honda et al. 2011; Honda et al. 2015), the domestic silk moth *Bombyx mori* (Hayashiya et al. 1965; Miyashita et al. 2016; Nguku et al. 2007), and the two-spotted field cricket *Gryllus bimaculatus* (Miyashita et al. 2016).

Royal jelly is composed of water (60-70%), proteins (8-18%), carbohydrates (7-18%), fatty acids and lipids (3-8%), and small amounts of vitamins and minerals (Kunugi and Ali 2019; Wytrychowski et al. 2013). Royal jelly’s variation in nutrient content depends in part on which honey bee species produced it and their geographical location, the timing of harvest, botanical source, storage conditions, pesticide exposure, and production methods (Al-Kahtani et al. 2020; Chaves et al. 2020; Kunugi and Ali 2019; Ma et al. 2021; Milone et al. 2021; Mokaya et al. 2020; Qi et al. 2020; Sano et al. 2004; Wytrychowski et al. 2013; Zheng et al. 2011). More than 80% of the protein in royal jelly is in the form of nine major royal jelly proteins (MRJP) (Albert and Klaudiny 2004; Klaudiny et al. 1994; Kunugi and Ali 2019; Xin et al. 2016). These MRJPs drive the specific physiological and life history changes that occur during queen development (Schmitzova et al. 1998). Dietary protein is important for providing amino acids required for growth and body size regulation (Han and Dingemanse 2017; Harrison et al. 2014; Reiffer et al. 2018), and royal jelly contains 17 free amino acids (Ahmad et al. 2020; Silici et al. 2009), 8 of which are essential for insects (lysine, valine, threonine, phenylalanine, leucine, isoleucine) (Cohen 2015). Protein is also important for insect egg production (Harrison et al. 2014; Joern and Behmer 1997; Roeder and Behmer 2014). While protein more strongly influences growth compared to carbohydrates when diets are provided *ad libitum* (Harrison et al. 2014; McDonald et al. 2011) the high sugar content (and thus high carbohydrate content) of royal jelly could still affect growth. Sugars are a phagostimulant that increases the quantity of food consumed (Buttstedt et al. 2016; Kunugi and Ali 2019), thus playing an indirect role in body size as individuals who consume more generally tend to grow bigger (Gutiérrez et al. 2020).

Insects are commonly used as model systems in nutritional ecology due to their short generation times, ease of rearing, and high reproductive output (Gutiérrez et al. 2020). These same life history traits make insect species amenable to mass rearing, such as those farmed for food and feed. Insects have been consumed across the globe for centuries, but insect farming and mass rearing facilities are relatively new to Europe and North America. Recently, North America has seen a burgeoning interest in entomophagy (Schrader et al. 2016), with 35 Canadian insect protein producers and companies offering insect-containing food products appearing in the last decade (NPC 2022). Crickets are economically important to these protein producing companies, as they are more desirable to consumers than larval-stage insects (e.g., mealworms and black soldier fly larvae) (Dossey et al. 2016; Reverberi 2020).

Advancements in diet technology and formulation could potentially reduce the costs associated with farming insects. Diet can have strong life history influences in insects (e.g. dietary protein and carbohydrates: Harrison et al. 2014, Cammack and Tomberlin 2017, Reifer et al. 2018, Kim et al. 2021; salts: Welti et al. 2019, Peterson et al. 2021; lipids: Blaul and Ruther 2011, Krabbe et al. 2019). Changes in diet formulations and/or the use of dietary supplements therefore have the potential to enhance yields of insect farms without additional labour. Miyashita et al. (2016) discovered that royal jelly feeding during development resulted in enlarged body size of both male and female crickets (*Gryllus bimaculatus*); this effect was dose-dependent and optimized at a 15% royal jelly diet. These results suggest that royal jelly (or its component MRJPs) is a dietary supplement that could enhance cricket farm yields. However, because effects of dietary royal jelly differ among invertebrate orders and species (Miyashita et al. 2016), mass-reared insect species regularly used by the industry may have different dietary requirements for optimal fitness than *Gryllus bimaculatus*, a species not used in this industry. The time is ripe to translate the effects of royal jelly to increasing the yield of a mass-reared insect model regularly used by the industry. We therefore determined the effects of royal jelly on male and female growth rate and adult body size of a mass reared insect to determine whether royal jelly could be used to grow farmed insects bigger and/or faster.

Here, we ran two experiments using a commercially reared cricket species, *Gryllodes sigillatus*, as our model organism. In Experiment 1 we asked: what is the minimum amount of royal jelly supplement required to elicit a significant morphological response? We then proceeded with a more in-depth investigation (Experiment 2) that determined the individual-level life history responses of *G. sigillatus* to royal jelly over time. We hypothesized that royal jelly would enhance growth in both males and females. Following Miyashita et al’s. (2016) findings on *Gryllus bimaculatus*, we predicted that both male and female *Gryllodes sigillatus* would grow larger when fed a control diet with a 15% royal jelly supplement compared to a control diet with no royal jelly supplement.

## 2.0 Materials and Methods

### 2.1 Cricket rearing

*Gryllodes sigillatus* eggs were hatched in a medium of peat moss inside an incubator (56 × 56 × 99 cm, Precision Scientific) maintained at ∼31.5-33°C and 60% RH. A 14:10 L:D cycle was maintained using a single LED light strip installed along the length of the incubator interior. These conditions were conserved for the duration of the experiments. Every other day, the peat moss and eggs were moistened with water to prevent desiccation and gently stirred to prevent mold growth. The eggs were monitored twice daily for emergence. Upon emergence, only those crickets that emerged within a 12-hour time frame were used for experiments. Newly emerged cricket nymphs were each placed into individual rearing containers made from 96.1 mL plastic condiment cups and lids. Twelve rearing containers were evenly spaced apart in a 4 × 3 grid on plastic cafeteria trays. Trays were assigned a new position inside the incubator every four days, and no tray was assigned the same position for the duration of the experiment. Diet treatments were randomly allocated for each tray. Each cricket was provided with a 14 mm wide polyethylene push-in cap for a food dish and a 0.75 uL PCR tube (lid removed) for a water vial which was stoppered with 38 mm wide saturated dental cotton wick (Healifty, China). Food and water were provided *ad libitum* and replaced every 3-4 days. After five weeks, all crickets were euthanized by freezing.

The standard diet (Earth’s Harvest Organic Cricket Grower) consisted of a mixture of 40.4% corn, 27.9% soybean meal, 15.0% fishmeal, and 1.70% vitamin and mineral mix (Nutri-mix Elite Grower DDG). Fresh 100% pure royal jelly (Planet Bee Honey Farm, Vernon, BC) was mixed with the standard diet to make treatment diets consisting of 10, 15, 20, and 30% w/w royal jelly. Diets were stored in a refrigerator at 4°C.

### 2.2 Measurements

Live individuals were placed into clear plastic Ziploc bags and photographed dorsally using a Dino-Lite Edge 3.0 Digital Microscope (Dunwell Tech, Inc, Torrance, CA, U.S.A) and DinoCapture 2.0 software (AnMo Electronics Corp., New Taipei City, Taiwan). The camera was calibrated for measurements prior to photographing specimens, and the subsequent photos were supplemented with a scale bar to accurately perform digital measurements. Head width (maximal distance between the outer edges of the eyes), pronotum width (maximal distance across the coronal width of the pronotum) and length (maximal distance down the sagittal length of the pronotum) were measured every seven days using ImageJ v.148 software (National Institutes of Health, Bethesda, MD, U.S.A). Eggs were photographed using the same materials and were manually counted in ImageJ using the Cell Counter plugin.

### 2.3 Experiment 1: Dose-response feeding curve

To determine the dose-dependent responses of *G. sigillatus* body size and mass to royal jelly supplements, and the probability of reaching adulthood when fed a royal jelly supplement, we reared *G. sigillatus* nymphs from hatch to adulthood on standard diets with a range of 0-30% royal jelly. Two separate trials were conducted. Upon emergence, cricket nymphs were weighed, photographed, and measured, and then placed into individual containers. For trial 1, diets consisting of 0%, 10%, 15%, and 20% royal jelly were fed to 25 crickets per diet (N = 100). In trial 2, diets consisting of 0%, 10%, 20%, and 30% were fed to 48 crickets per diet (N = 192) for a total of 292 crickets in Experiment 1 collected across the two trials. Body size and weight measurements were performed 35 days after hatch.

### 2.4 Experiment 2 –15% Royal Jelly Supplement

We tested how adding a 15% royal jelly supplement to the standard diet affected the growth and egg production of *G. sigillatus*. Two separate trials were conducted, and in each trial 96 crickets were fed either the standard diet with no royal jelly supplement (control, N = 48) or the standard diet with a 15% royal jelly supplement (N = 48) for a total of 192 crickets in Experiment 2. Body mass was measured weekly starting seven days after hatch. Body size and mass measurements were performed weekly for 35 days after hatch, and the crickets were then frozen. We performed post-hoc measurements on abdomen length (maximal distance down the sagittal length of the abdomen) and the number of eggs produced per cricket (N=15 basal, N=28 15% royal jelly).

Abdomens were opened with a ventral longitudinal incision and a smaller perpendicular horizontal incision near the anal opening. The digestive tract was removed, and all eggs were flushed with water out from the abdominal cavity. An image of the eggs was captured, and the number of eggs was counted manually.

### 2.5 Data analysis

All statistical analyses were conducted using JMP Pro 15 software (SAS Institute Inc., 2014). Normality of the data was confirmed through residual analyses and Shapiro-Wilk goodness-of-fit tests for each variable (*P* > 0.05). We used the false discovery rate B-Y method (FDR_BY_) to adjust alpha for each statistical test (Benjamini and Yekutieli 2001). Trial was included as a random effect in all analyses and was never statistically significant (*P* > 0.05).

#### Data analysis – Experiment 1: Dose-response feeding curve

Data from Experiment 1 (0%, 10%, 15%, 20%, and 30% royal jelly supplements) and 2 (0% and 15% royal jelly supplements) were pooled to increase sample size for a total of 464 crickets (N = 169, 73, 121, 73, and 48 for the 0, 10, 15, 20, and 30% royal jelly diets, respectively). In total, 304 crickets survived and 280 reached adulthood (see Appendix A for survival distribution). Crickets that did not reach adulthood by the end of the experiment were removed from the analyses. To explore how royal jelly supplements affected the probability that individual crickets would reach adulthood we used logistic regression. The categorical dependent variable was whether the individual was a juvenile or an adult at 35 days post imaginal moult, and the independent variable was the percentage of royal jelly (w/w) in the diet.

To determine whether royal jelly concentration impacted adult mass and body size we used linear mixed models (LMM). If significant linear and quadratic regressions were found for a variable, AICc values were used to select the best fit. Diet (0%, 10%, 15%, 20%, 30% royal jelly) and sex (female or male) were included as fixed effects, and experiment trial (Experiment 1: trials 1 and 2; Experiment 2: trials 3 and 4) was included as a random effect.

#### Data analysis - Experiment 2: 15% Royal jelly supplement

161 of 192 total crickets reached adulthood in Experiment 2. Crickets that did not reach adulthood by the end of the experiment were removed from the analysis. Of the surviving 161 crickets, 45 males and 33 females were fed the basal diet, and 36 males and 47 females were fed the royal jelly diet. A chi-squared test was performed to test if these ratios were significantly different from expected. Principal component analysis was used to extract orthogonal vectors from head width, pronotum width, and pronotum length at adulthood to quantify adult body size into a single measurement. The first principal component (PC1) explained 92.12% of the variation in size (eigenvalue = 2.76); all size measures loaded equally on PC1. We fit a LMM to determine whether diet treatment influenced mass, body size (PC1), head width, pronotum width, pronotum length, and abdomen length. In all analyses, diet [standard diet only (control) or standard diet with 15% royal jelly supplement], age (days since hatch), sex (male or female), and all possible interactions were included as fixed effects. Given the repeated aspects of the experiment, both cricket identification number and trial were included as random effects. We used a linear model (LM) to determine whether royal jelly impacted day-of-eclosion adult mass, body size, (PC1), head width, pronotum width, pronotum length, and abdomen length. Diet, sex, and the interaction between diet and sex were included as fixed effects, and trial was included as a random effect. A separate LM was used to determine whether royal jelly impacted the number of eggs at adulthood with diet as the only effect in the model because only females produce eggs and eggs were dissected only in trial 1. Pairwise comparisons for the LM were developed using Tukey’s HSD.

## 3.0 Results

### Dose-response feeding curve

The odds ratio of crickets reaching adulthood after 35 days was 19.2 higher for crickets fed royal jelly (*X*^*2*^ = 14.27; *P* < 0.001; R^2^ = 0.12). Significant linear and quadratic relationships were found between royal jelly concentration and female mass and abdomen length (Figure 1; Table 1), and the quadratic relationships were both a better fit than linear based on lower AICc values. The quadratic relationships between royal jelly and female mass and abdomen length predicted maximized responses when fed 16.95% and 17.60% royal jelly, respectively.

**Figure 1.**
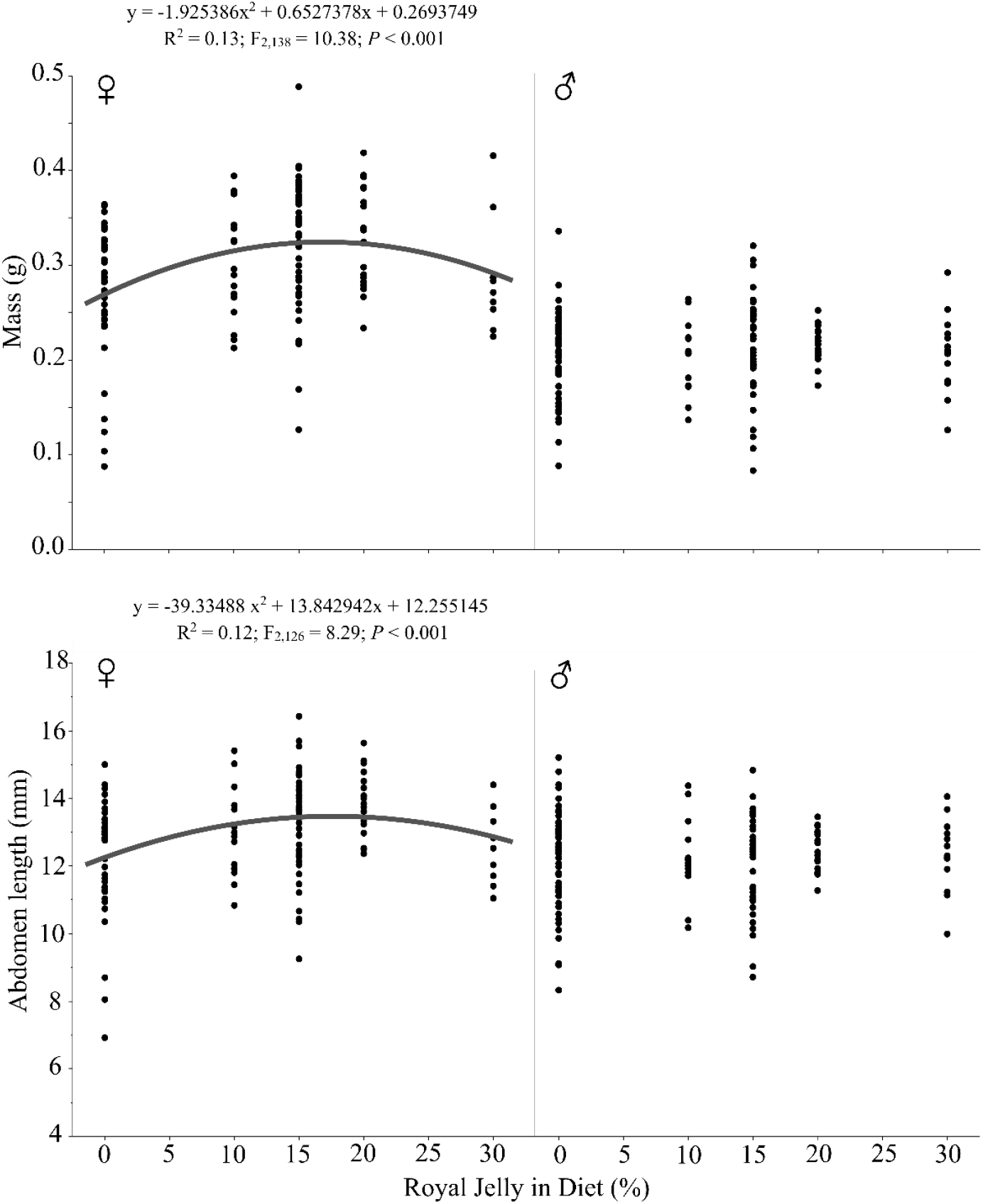
Royal jelly dose-response regressions of *Gryllodes sigillatus*. Female mass and abdomen length are explained by significant quadratic relationships.

**Table 1.** Summary statistics from the linear mixed model of 35-day old adult *Gryllodes sigillatus* measurements from Experiment 1: Dose-response feeding curve.

| Model | R<sup>2</sup> | Term | DF | F | P |
| --- | --- | --- | --- | --- | --- |
| Mass | 0.53 | Diet | 1,272.60 | 15.68 | **<0.001** |
|  |  | Diet<sup>2</sup> | 1,274 | 5.17 | **0.030** |
|  |  | Sex | 1,271.20 | 215.57 | **<0.001** |
|  |  | Diet*Sex | 1,271.60 | 7.43 | **0.011** |
|  |  | Diet<sup>2</sup>*Sex | 1,271.50 | 4.37 | **0.038** |
| Body Size (PC1) | 0.49 | Diet | 1,270.70 | 0.90 | 0.57 |
|  |  | Diet<sup>2</sup> | 1,271.90 | 0.038 | 0.85 |
|  |  | Sex | 1,270.10 | 167.45 | **<0.001** |
|  |  | Diet*Sex | 1,270.20 | 0.92 | 0.57 |
|  |  | Diet<sup>2</sup>*Sex | 1,270.10 | 0.31 | 0.72 |
| Abdomen Length | 0.26 | Diet | 1,257.60 | 8.94 | **0.0076** |
|  |  | Diet<sup>2</sup> | 1,258.80 | 4.42 | **0.037** |
|  |  | Sex | 1,256.20 | 21.36 | **<0.001** |
|  |  | Diet*Sex | 1,256.50 | 7.57 | **0.011** |
|  |  | Diet<sup>2</sup>*Sex | 1,256.40 | 5.08 | **0.031** |

### 15% Royal Jelly Supplement

The ratio of crickets that survived and developed into males and females was significantly different than expected (*X*^*2*^ (1) = 3.89; *P* = 0.049). However, this result was made less clear after pooling the 0% and 15% data across the two experiments and running the same chi-squared test: there was no difference between expected and observed ratios (*X*^*2*^ (1) = 2,34; *P* = 0.13). The 15% royal jelly supplement significantly affected *G. sigillatus* mass (*P* < 0.001) and pronotum length (*P =* 0.025), but males and females were affected differently (Table 2). Females grew heavier (*P* = 0.031) when fed a royal jelly supplement, but the royal jelly supplement did not significantly affect overall body size as encapsulated by PC1 (*P* = 0.07; Table 2). Over time females increased mass, grew larger bodies (PC1), and grew longer pronotums more rapidly when they were fed 15% royal jelly supplements compared to the standard diet without supplements (*P* < 0.05; Table 2; Figure 2). At day 35, adult females fed royal jelly were 30% heavier on average, had significantly longer abdomens, and had significantly higher fecundity (66.6% more eggs) than females fed the standard diet (Figure 3; Tables 3, 4). Male mass and body size were unaffected by royal jelly both over time and at the end of the experiment (Figures 2, 3). Trial was never a significant source of variation (*P* > 0.05).

**Table 2.** Summary statistics from the repeated measures linear mixed model of *Gryllodes sigillatus* measurements from Experiment 2: Direct test of 15% royal jelly

| Model | R<sup>2</sup> | Term | DF | F | P |
| --- | --- | --- | --- | --- | --- |
| Mass | 0.84 | Diet | 1, 153.3 | 11.74 | **<0.001** |
|  |  | Sex | 1, 153.3 | 37.23 | **<0.001** |
|  |  | Age | 1, 712.1 | 3533.24 | **<0.001** |
|  |  | Diet*Sex | 1, 153.4 | 4.75 | **0.031** |
|  |  | Diet*Age | 1, 732.9 | 16.71 | **<0.001** |
|  |  | Sex*Age | 1, 732.9 | 102.34 | **<0.001** |
|  |  | Diet*Sex*Age | 1, 732.9 | 12.04 | **<0.001** |
| Body Size (PC1) | 0.96 | Diet | 1, 156 | 4.08 | 0.063 |
|  |  | Sex | 1, 156 | 9.33 | **0.0062** |
|  |  | Age | 1, 797.5 | 17054.58 | **<0.001** |
|  |  | Diet*Sex | 1, 156.1 | 3.58 | 0.070 |
|  |  | Diet*Age | 1, 797.5 | 1.76 | 0.19 |
|  |  | Sex*Age | 1, 797.5 | 86.65 | **<0.001** |
|  |  | Diet*Sex*Age | 1, 797.5 | 4.86 | **0.049** |
| Head width | 0.96 | Diet | 1, 156 | 2.68 | 0.10 |
|  |  | Sex | 1, 156 | 4.75 | **0.031** |
|  |  | Age | 1, 797.4 | 19372.19 | **<0.001** |
|  |  | Diet*Sex | 1, 156.1 | 2.94 | 0.088 |
|  |  | Diet*Age | 1, 797.4 | 0.63 | 0.43 |
|  |  | Sex*Age | 1, 797.4 | 73.49 | **<0.001** |
|  |  | Diet*Sex*Age | 1, 797.4 | 2.80 | 0.094 |
| Pronotum width | 0.96 | Diet | 1, 156.1 | 4.14 | 0.06 |
|  |  | Sex | 1, 156 | 8.71 | **0.0085** |
|  |  | Age | 1, 797.5 | 17363.46 | **<0.001** |
|  |  | Diet*Sex | 1, 156.2 | 3.52 | 0.073 |
|  |  | Diet*Age | 1, 797.5 | 2.79 | 0.096 |
|  |  | Sex*Age | 1, 797.5 | 80.90 | **<0.001** |
|  |  | Diet*Sex*Age | 1, 797.5 | 4.25 | 0.061 |
| Pronotum length | 0.94 | Diet | 1, 155.9 | 5.70 | **0.025** |
|  |  | Sex | 1, 155.9 | 17.43 | **<0.001** |
|  |  | Age | 1, 797.5 | 11711.02 | **<0.001** |
|  |  | Diet*Sex | 1, 156 | 4.18 | 0.050 |
|  |  | Diet*Age | 1, 797.5 | 1.99 | 0.16 |
|  |  | Sex*Age | 1, 797.5 | 92.92 | **<0.001** |
|  |  | Diet*Sex*Age | 1, 797.5 | 7.58 | **0.011** |

**Figure 2.**
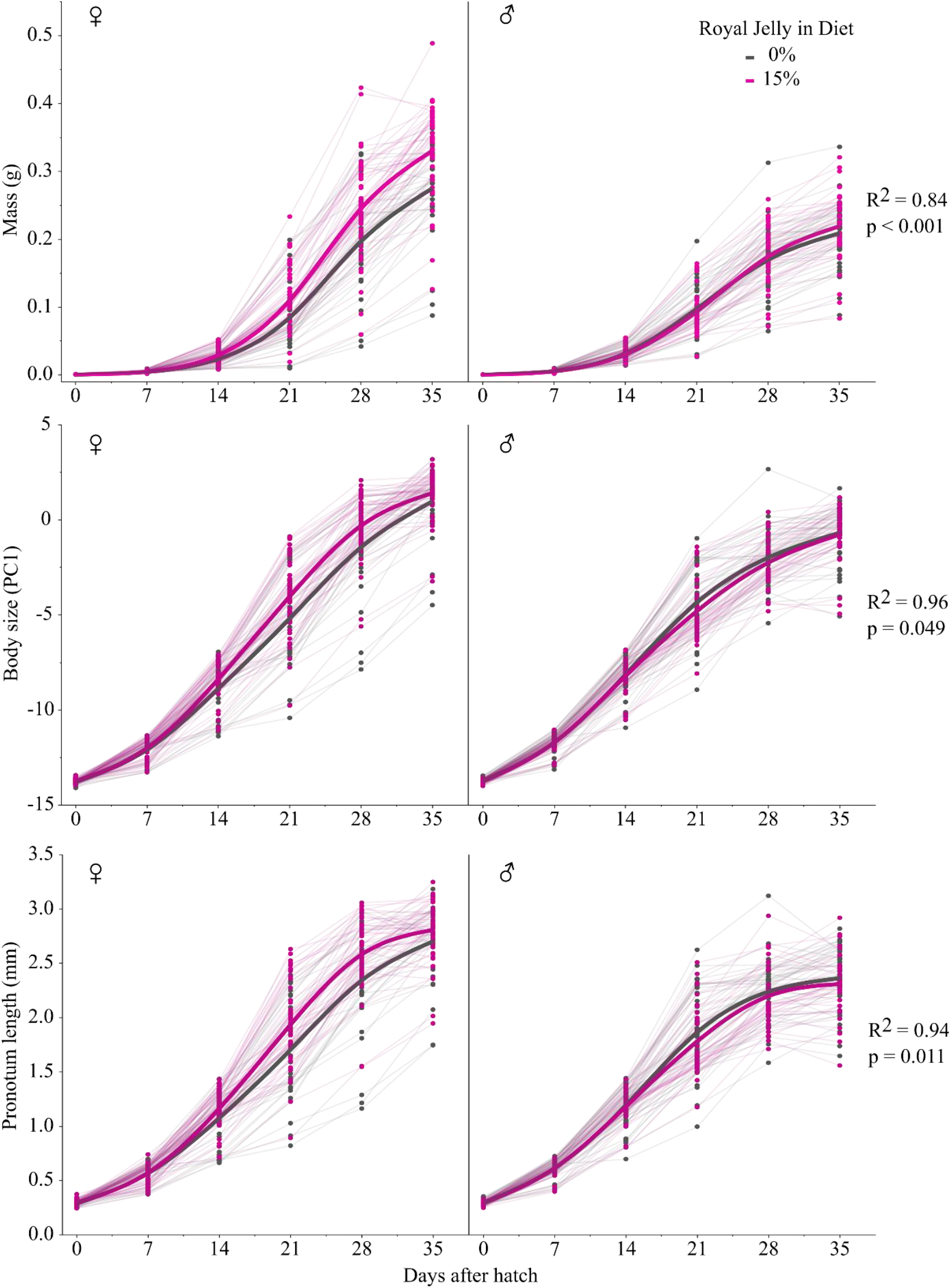
Lifetime development of mass, body size (PC1), and pronotum length over age of *Gryllodes sigillatus* fed either a basal diet or a diet with 15% royal jelly. Smoothed thick lines connect mean values at each timepoint, and solid circles represent individual crickets and are connected by thin lines over time.

**Figure 3.**
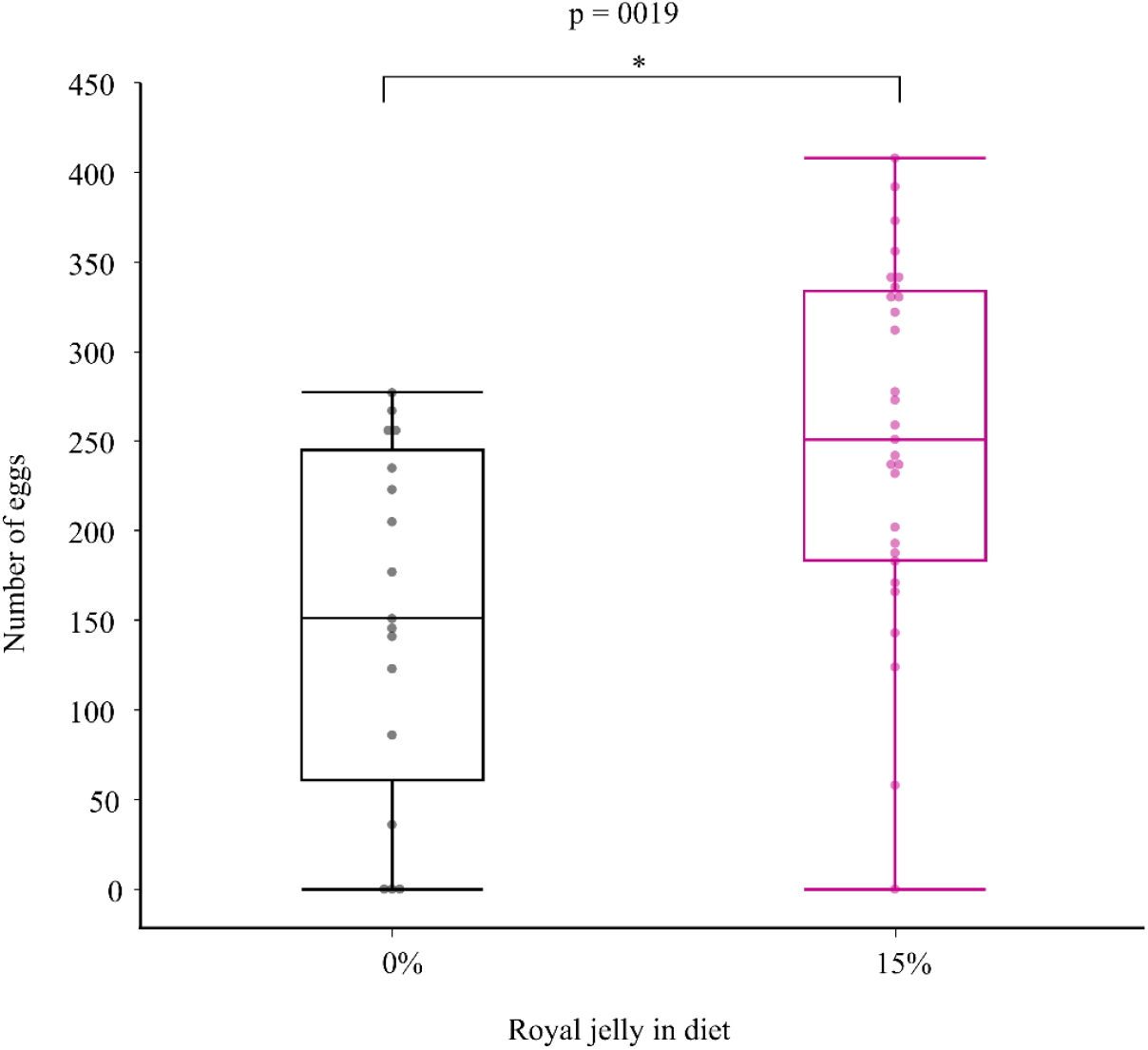
Number of eggs dissected from 35-day old adult *Gryllodes sigillatus* females fed basal and 15% royal jelly diets. Auto jitter was applied to the data points in JMP Pro Graph Builder.

**Table 3.**
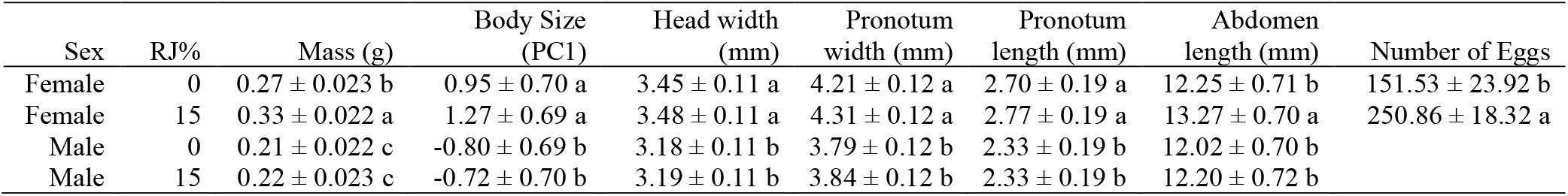
Least square means ± standard error of morphology and life history measurements of 35-day old adult *Gryllodes sigillatus* from Experiment 2: Direct test of 15% royal jelly.

| Sex | RJ% | Mass (g) | Body Size<br>(PC1) | Head width<br>(mm) | Pronotum<br>width (mm) | Pronotum<br>length (mm) | Abdomen<br>length (mm) | Number of Eggs |
| --- | --- | --- | --- | --- | --- | --- | --- | --- |
| Female | 0 | 0.27 ± 0.023 b | 0.95 ± 0.70 a | 3.45 ± 0.11 a | 4.21 ± 0.12 a | 2.70 ± 0.19 a | 12.25 ± 0.71 b | 151.53 ± 23.92 b |
| Female | 15 | 0.33 ± 0.022 a | 1.27 ± 0.69 a | 3.48 ± 0.11 a | 4.31 ± 0.12 a | 2.77 ± 0.19 a | 13.27 ± 0.70 a | 250.86 ± 18.32 a |
| Male | 0 | 0.21 ± 0.022 c | -0.80 ± 0.69 b | 3.18 ± 0.11 b | 3.79 ± 0.12 b | 2.33 ± 0.19 b | 12.02 ± 0.70 b | |
| Male | 15 | 0.22 ± 0.023 c | -0.72 ± 0.70 b | 3.19 ± 0.11 b | 3.84 ± 0.12 b | 2.33 ± 0.19 b | 12.20 ± 0.72 b | |

**Table 4.** Summary statistics from the linear model of 35-day old adult *Gryllodes sigillatus* measurements from Experiment 2: Direct test of 15% royal jelly.

| Model | R<sup>2</sup> | Term | DF | F | P |
| --- | --- | --- | --- | --- | --- |
| Mass | 0.51 | Diet | 1, 156 | 15.45 | **<0.001** |
|  |  | Sex | 1, 156 | 103.95 | **<0.001** |
|  |  | Diet*Sex | 1, 156.2 | 4.25 | **0.041** |
| Body Size (PC1) | 0.45 | Diet | 1, 156 | 0.97 | 0.49 |
|  |  | Sex | 1, 156 | 79.97 | **<0.001** |
|  |  | Diet*Sex | 1, 156.1 | 0.31 | 0.58 |
| Abdomen Length | 0.25 | Diet | 1, 142.1 | 6.20 | **0.021** |
|  |  | Sex | 1, 142 | 7.24 | **0.021** |
|  |  | Diet*Sex | 1, 142.1 | 3.01 | 0.085 |
| Number of Eggs | 0.18 | Diet | 1, 44 | 10.87 | **0.0019** |

## 4.0 Discussion and Conclusion

Evidence from several insect orders and nematodes suggests that honey bee royal jelly, when administered orally through a diet, can influence life history trait development. Key similarities among insects include increased body size and mass (*D. melanogaster*, Arruda et al. 2017; Kamakura 2011; Shorter et al. 2015; *B. mori*, Hayashiya et al. 1965; Miyashita et al. 2016; Nguku et al. 2007; *Gryllus bimaculatus*, Miyashita et al. 2016), and increased fecundity (*D. melanogaster*, Kamakura 2011; Kayashima et al. 2012; *B. mori*, Miyashita et al. 2016). Despite this body of past work, the present study is the first to our knowledge that has tested the effects in an insect species mass reared for food and feed. We found a sex-specific effect of royal jelly in a hemimetabolous insect (*Gryllodes sigillatus*): females grew heavier faster and developed more eggs when fed royal jelly, but males were unaffected. These results complicate our understanding of the effects of royal jelly in insects, as they complement but also contrast with Miyashita et al.’s (2016) finding that both male and female *Gryllus bimaculatus* (a cricket in the same family as *G. sigillatus*) grew larger when fed royal jelly. In the same study, sex-specific differences like those we observed here were found in *B. mori* (a lepidopteran); female moth pupae and adults grew larger and developed more eggs when fed royal jelly. Miyashita et al. (2016) suggested that the response differences to royal jelly observed in *B. mori* and *Gryllus bimaculatus* were a result of different developmental processes and mechanisms that respond to royal jelly between holo- and hemimetabolous insects. While the theory of insects with different development life histories responding to diet differently is not new (Bernays 1986; Bertram et al. 2021; Neiro 2020; Thompson 2019), our results suggest that the effects of royal jelly can be sex-specific in Orthoptera. Thus, distantly-related holometabolous and hemimetabolous insects can respond similarly to dietary royal jelly, while two relatively closely related hemimetabolous orthopterans can respond differently. The signaling mechanisms activated by royal jelly molecules may thus be conserved between holometabolous and hemimetabolous insects, but the ultimate effects of royal jelly might be tied to other life history traits or the dietary or environmental context in which it is experienced by an insect. Miyashita et al. (2016) concluded that royal jelly affected cricket size via upregulation of food consumption, and while this would have been an excellent addition to our study, limitations brought on by the COVID-19 pandemic severely restricted access to resources necessary to accurately measure food consumption.

Sex-specific life-history differences in response to diet are well documented in insects, especially in crickets. Protein intake is important for optimizing fecundity and egg production in females, and carbohydrate intake regulates calling effort in males (*Teleogryllus oceanicus*: Ng et al. 2018; *T. commodus*: Maklakov et al. 2008; *Gryllus veletis*: Harrison et al. 2014; *Gryllus assimilis*: Reifer at al. 2018). Houslay et al. (2015) provide a stimulating discussion on the difference in resource acquisition between females and males; females have a straightforward process of gathering resources and converting them into eggs, whereas males exhibit acquisition-related plasticity and must choose how to allocate invested resources (calling effort, competition, etc.). The investment strategy of male *G. sigillatus* can explain the differences between sexes in our experiment; males exhibit acquisition-related plasticity in calling behaviour (Bertram et al. 2011; Houslay et al. 2015), and so compared to females who become fecund by gathering nutrients and converting those nutrients into eggs, males experience the resource-intensive additional burden of producing a spermatophylax and will conserve resources and energy until they perceive a competitor (Mallard and Barnard 2003, 2004). Investment strategy and resource acquisition can also explain the response differences to royal jelly between *G. sigillatus* and *Gryllus bimaculatus*. Both species increase the number of sperm transferred in the presence of apparent competitors; male *G. sigillatus* increase significantly more sperm with a conspecific competitor whereas *Gryllus bimaculatus* increase sperm regardless of the competitor species (Mallard and Barnard 2003). In a group-environment rearing protocol used by Miyashita et al. (2016), it is no surprise that both male and female *Gryllus bimaculatus* increased body size on a royal jelly (higher quality) diet; males were susceptible to competition, and likely increased their investment by consuming more food, and females grow heavier under higher population density (EL-Damanhouri 2011). Under the isolated and individual-rearing conditions used here, we suspect that *G. sigillatus* were less able to detect competition and thus there was no competitor cue to increase investment in body size. This hypothesis could be tested by repeating our experiment in a controlled group-rearing and/or active farm to test whether royal jelly also influences body size of male *G. sigillatus* in a group environment. If *G. sigillatus* were fed royal jelly in a mass rearing environment, with the constant presence of conspecific competition, males may demonstrate a response of increased body size regulated by increased investment.

In addition to farm-scale trials, future research should also explore how royal jelly, either fresh or manufactured, impacts male and female growth rate and body size over generational time. Lab populations of *Plutella xylostella* (Lepidoptera: Plutellidae) reared on high-carbohydrate diets initially responded with reduced fitness, however over 350 generations they evolved the ability to reduce fat storage (Warbrick-Smith et al. 2009). Hall et al. (2008) reared Australian ground cricket (*Pteronemobius sp*.) for seven generations on low- and high-quality diets, and while individuals at the beginning of the experiment grew heavier on the high-quality diet, after seven generations crickets grew significantly heavier on the low-quality diet treatment. Therefore, single generation diet changes may be misleading in their long-term effects but are still valuable for demonstrating the potential power that dietary supplements may have. Further research should test royal jelly across multiple generations of *G. sigillatus* to determine how it may impact overall farm yield, and whether these sorts of supplements will be financially viable for use in entomophagical farming.

Our study is a useful contrast to Miyashita et al’s (2016) for highlighting species-specific nutritional requirements to optimize fitness. Our study also represents an important step toward application of nutritional supplements in cricket farming. *Gryllodes sigillatus* is an important economical cricket species currently farmed for food and feed in North America. Small and medium-scale cricket farms often use whatever feed is safe, available, and consumed by the insects – if the crickets eat, grow, and reproduce, the feed is deemed acceptable (Entomo Farms, personal communication). This keeps production costs low but can lead to lower harvest yields and variable nutrient quality of the food products (Weru et al. 2021). Our results contribute to a growing body of literature that demonstrates the strong potential of using royal jelly as a dietary supplement to optimize the mass-rearing of insect species. Although royal jelly is expensive, difficult to produce in large quantities (Ramanathan et al. 2018), and quick to spoil in storage (Tarantilis et al. 2012), research is underway into improving the mass production of natural and artificial royal jelly proteins responsible for enhancing body size and growth (Cao et al. 2016; Ma et al. 2021; Wang et al. 2020). Synthesizing royal jelly into mass reared insect diets must be cost effective but also maintain or increase the gustatory appeal of the diet. Royal jelly is rich in free amino acids (Guo et al. 2021), and female *G. sigillatus* feeding time increases with spermatophylax amino acid concentration (Warwick et al. 2009) thus increasing the total amount of ejaculate transferred to females. Diet can have a strong influence on male *G. sigillatus* fitness (Duffield et al. 2020), and male mating success is maximized on diets slightly higher in protein than carbohydrates (Rapkin et al. 2017). Therefore, it would be valuable for future studies to test how dietary royal jelly influences spermatophylax composition, feeding time, and overall mating success in *G. sigillatus*.

## Acknowledgements

We thank Olivia Gagnon and Kylie Barwise for laboratory assistance. Thank you to Entomo Farms for providing specimens and diet.

## Appendices

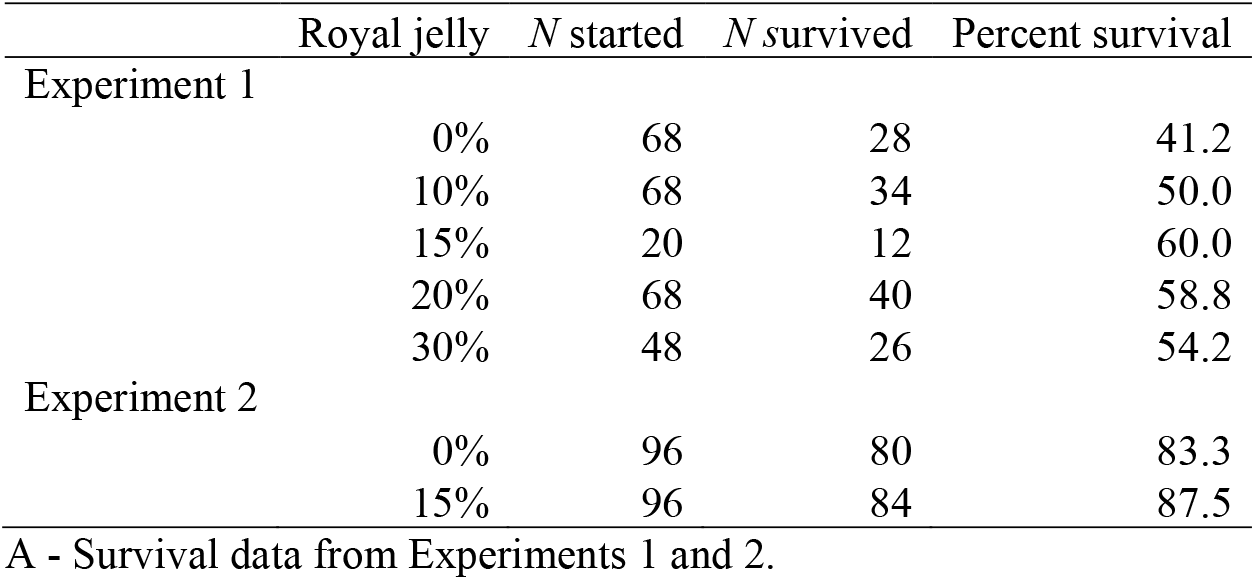

A - Survival data from Experiments 1 and 2.
| Model | R<sup>2</sup> | Term | DF | F | P |
| --- | --- | --- | --- | --- | --- |
| Head width | 0.41 | Diet | 1,270.90 | 0.037 | 0.85 |
|  |  | Diet<sup>2</sup> | 1,272.40 | 0.28 | 0.79 |
|  |  | Sex | 1,270.10 | 115.55 | **<0.001** |
|  |  | Diet*Sex | 1,270.30 | 0.66 | 0.79 |
|  |  | Diet<sup>2</sup>*Sex | 1,270.20 | 0.23 | 0.79 |
| Pronotum width | 0.41 | Diet | 1,271.20 | 2.86 | 0.23 |
|  |  | Diet<sup>2</sup> | 1,272.80 | 0.92 | 0.45 |
|  |  | Sex | 1,270.10 | 133.83 | **<0.001** |
|  |  | Diet*Sex | 1,270.40 | 0.85 | 0.45 |
|  |  | Diet<sup>2</sup>*Sex | 1,270.30 | 0.32 | 0.57 |
| Pronotum length | 0.62 | Diet | 1,270.30 | 0.57 | 0.73 |
|  |  | Diet<sup>2</sup> | 1,271 | 0.0011 | 0.97 |
|  |  | Sex | 1,270 | 233.20 | **<0.001** |
|  |  | Diet*Sex | 1,270.10 | 1.068 | 0.73 |
|  |  | Diet<sup>2</sup>*Sex | 1,270.10 | 0.31 | 0.73 |

B - Summary statistics from the linear mixed model of 35-day old *Gryllodes sigillatus* measurements from Experiment 1: Dose-response feeding curve for measurements included and combined into principal component 1 (body size).

| Model | R<sup>2</sup> | Term | DF | F | P |
| --- | --- | --- | --- | --- | --- |
| Head width | 0.36 | Diet | 1, 156 | 0.28 | 0.85 |
|  |  | Sex | 1, 156 | 54.87 | **<0.001** |
|  |  | Diet*Sex | 1, 156.1 | 0.035 | 0.85 |
| Pronotum width | 0.36 | Diet | 1, 156 | 1.81 | 0.27 |
|  |  | Sex | 1, 156 | 64.46 | **<0.001** |
|  |  | Diet*Sex | 1, 156.2 | 0.26 | 0.61 |
| Pronotum length | 0.60 | Diet | 1, 156 | 0.92 | 0.34 |
|  |  | Sex | 1, 156 | 121.4 | **<0.001** |
|  |  | Diet*Sex | 1, 156 | 1.055 | 0.34 |

C - Summary statistics from the linear model of 35-day old adult *Gryllodes sigillatus* measurements from Experiment 2: Direct test of 15% royal jelly for measurements included and combined into principal component 1 (body size).

